# Engineering a Covalent Linkage into a Dimeric *De Novo* Enzyme Reveals a Novel Life-Sustaining Mechanism

**DOI:** 10.64898/2025.12.09.693225

**Authors:** Guanyu Liao, Sha Tao, Mina Nagahara, Kodai Kurihara, Koji Umezawa, Ryoichi Arai, Michael H. Hecht

## Abstract

Designing novel proteins that share no homology with natural sequences, but which nonetheless provide life sustaining functions, is an important goal for synthetic biology. Towards this goal, we previously reported Syn-F4, the first *de novo* enzyme capable of catalyzing a life-sustaining reaction, both *in vitro* and *in vivo*. Syn-F4 catalyzes hydrolysis of the siderophore, ferric enterobactin, thereby releasing iron and enabling growth in iron-limited media of an otherwise inviable Δ*fes* strain of *Escherichia coli*. Although Syn-F4 provides a direct enzymatic replacement of the natural ferric enterobactin esterase encoded by Fes, it has a dramatically different structure and enzymatic mechanism than the natural Fes enzyme. The novel Syn-F4 enzyme forms a 4-helix bundle, comprising a homodimer of two α-helical hairpins. Here we describe the engineering of a covalent peptide linkage into the homodimer to generate a single chain 4-helix bundle. As expected, the resulting linked protein (Syn-F4-Link) also rescued Δ*fes* cells in iron-limited media. Moreover, X-ray crystallography revealed a 3D structure similar to the parental homodimer. Surprisingly, however, the linked protein was *not* enzymatically active. Instead, Syn-F4-Link rescues Δ*fes* cells by upregulating biosynthesis of the enterobactin siderophore thereby enabling assimilation of sufficient iron to sustain cell growth. These findings demonstrate that two very similar *de novo* proteins can sustain cell growth using dramatically different biological mechanisms.

## INTRODUCTION

Natural proteins perform a wide range of functions necessary to sustain the growth of living organisms. Some proteins are structural, others are catalytic, and yet others perform regulatory functions necessary to modulate cell growth. However, despite the enormous diversity of functions, structures, and amino acid sequences observed in biology, the total collection of proteins in all biological systems on earth (past and present) represents a miniscule fraction of what might be possible.^1–3^ Indeed, the burgeoning field of *de novo* protein design – recognized recently by a Nobel Prize – demonstrates that it is now possible to devise entirely new sequences that fold into novel 3-dimensional structures not seen in nature.^4–7^ Recent advances in protein design have also demonstrated the possibility of creating proteins with a range of novel functions.^8–14^

Our previous work in protein design focused on the synthesis of large combinatorial libraries of novel sequences, followed by screens and selections aimed at discovering non-natural proteins with new functions.^1,15,16^ We focused not only on proteins that function in a test tube, but more importantly, on novel proteins that perform essential functions in living cells. For example, we discovered novel proteins that enable the growth of *E. coli* in concentrations of copper that would otherwise be toxic.^17^ We also discovered novel proteins that rescue nutritional auxotrophs in which an essential gene is deleted.^18^ For example, the deletion of *serB* (which encodes phosphoserine phosphatase) or *gltA* (which encodes citrate synthase) prevent the growth of *E. coli* in media lacking serine or glutamic acid, respectively. In both these cases, we isolated *de novo* proteins that rescue these deletions by altering gene regulation to increase the expression of alternative endogenous enzymes that have promiscuous activities, which if expressed at high levels, can rescue the auxotrophic strain.^19,20^

Novel proteins might also function *in vivo* by directly providing an essential enzymatic activity. For example, the growth of *E. coli* in iron-limited media depends on the endogenous enzyme Ferric Enterobactin Esterase (Fes), which degrades the siderophore Ferric Enterobactin (Fe(III)-Ent) to release iron and make it available for cell growth.^21–23^ Mutant cells in which this enzyme is deleted (Δ*fes)* fail to grow in iron poor media.^24^ Several years ago, we described the isolation of a binary patterned 4-helix bundle *de novo* protein, called Syn-F4, which rescues Δ*fes* cells.^25^ Purification of Syn-F4 confirmed that the novel protein is catalytically active as an enterobactin esterase, and moreover, that it is stereospecific.^25^ Recently, we reported the crystal structure of Syn-F4, demonstrating that it is a homodimeric 4-helix bundle with the C-terminus of one chain in close proximity to the N-terminus of the second chain.^26^ Model building and extensive mutagenesis studies suggested the catalytic site is at the center of the 4-helix bundle comprising residues from both polypeptide chains.^26^

The homodimeric 4-helix bundle structure of Syn-F4 is entirely different from the larger and more complex structure of the natural Fes enzyme. Moreover, the *de novo* protein catalyzes hydrolysis of its Fe(III)-Ent substrate using a mechanism that differs from that of the naturally evolved enzyme: whereas the natural enzyme is a classic serine hydrolase with a catalytic triad comprising serine, histidine, and aspartic acid, Syn-F4 contains no serine in its entire sequence and uses water as a nucleophile to hydrolyze the enterobactin substrate.^26^ The fact that the sequence, structure, and catalytic mechanism of Syn-F4 differ so dramatically from those of the natural Fes protein provide compelling evidence that life-essential biological challenges can be solved in ways that are very different from those selected by evolution.^26^

Because the termini of the two chains are close to one another in the crystal structure of the Syn-F4 homodimer, we surmised that linking these chains into a single double-length polypeptide would diminish the entropic cost of dimerization and thereby produce a more stable and/or more active enzyme. Here we describe construction of the covalent dimer and report that its crystal structure, as expected, is similar to that of the non-covalent dimer. Surprisingly, however, the enzymatic properties of these very similar proteins are dramatically different: the linked protein *cannot* catalyze the hydrolysis of Fe(III)-Ent. Consequently, although both the noncovalent dimer (Syn-F4) and the newly constructed covalent dimer (Syn-F4-Link) rescue Δ*fes* cells *in vivo,* they do so using distinctly different biological mechanisms.

## RESULTS

### Creation of a covalent dimer of Syn-F4 by inserting a flexible linker

Previously, we determined the crystal structure of Syn-F4 and found that it forms a homodimeric 4-helix bundle, with the N- and C-termini of both chains occurring at the same end of the bundle (*syn* topology, **SI Figure 1**).^26^ Based on this structure, we assumed that linking the two chains with a short flexible peptide would produce a covalently linked homodimer with a structure and properties similar to the original non-covalent dimer of Syn-F4. Toward this goal, we constructed a gene encoding a protein in which the two chains are joined by the flexible decapeptide, Gly-Gly-Gly-Gly-Ser-Gly-Gly-Gly-Gly-Ser (**Figure 1A**). We also made two mutations on the linked version of Syn-F4 at K4T and K116T corresponding to the previous mutation K4T on both chains of Syn-F4, which had been incorporated to enhance crystallization. We named this covalent dimer Syn-F4-Link K4Tx2, or Syn-F4-Link in brief.

**Figure 1.**
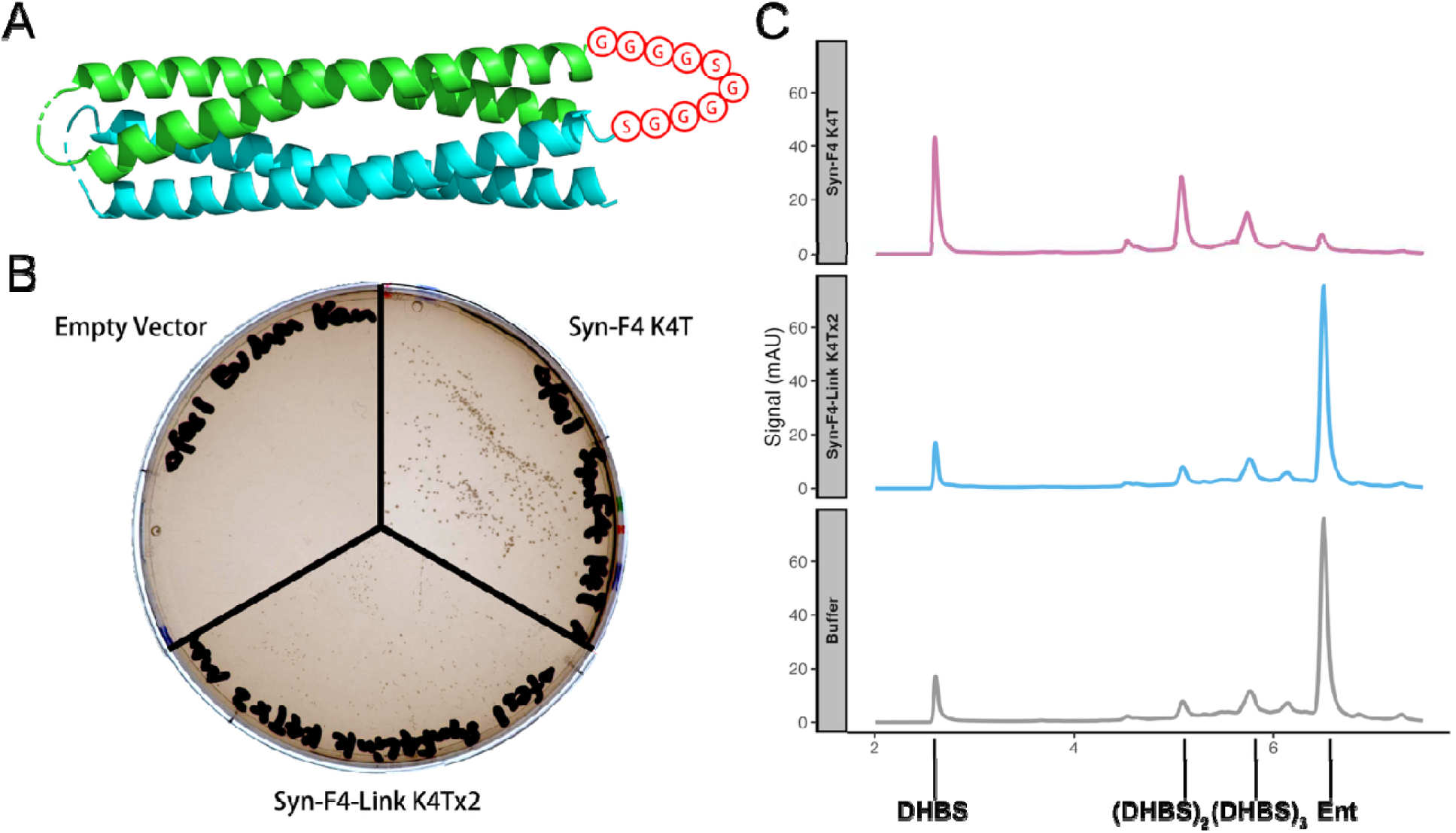
Syn-F4-Link rescues **Δ***fes* but cannot hydrolyze Fe(III)-Ent *in vitro*. (A) Design of Syn-F4-Link. The linker residues are shown in red circles. (B) Rescue of *E. coli* growth by Syn-F4, Syn-F4-Link, with empty vector as a negative control. (C) Hydrolysis of Fe(III)-Ent by purified Syn-F4, Syn-F4-Link (and buffer as negative control) monitored by reverse phase HPLC. Enterobactin (Ent), and its hydrolysis products (DHBS, (DHBS)_2_ and (DHBS)_3_) are labeled.

### Syn-F4-Link and the non-covalent Syn-F4 homodimer have similar structures, yet slightly different active site conformations

We solved the crystal structure of Syn-F4-Link and found it to be nearly identical to that of the original Syn-F4, with C-alpha RMSD = 0.45 Å (**Figure 2A, B**). The catalytic residues for Fe(III)-Ent hydrolysis have an all-atom RMSD of 0.78 Å, with Glu26 accounting for the slightly larger RMSD among the active site residues (RMSD = 1.0 Å) (**Figure 2C**). This shift in the orientation of Glu26 disables the H-bonding between Glu26, His74, and water, which was seen in the original Syn-F4 structure (**Figure 2D**). This alteration may prevent formation of the catalytic dyad, and therefore hamper the activity for Fe(III)-Ent hydrolysis.

**Figure 2.**
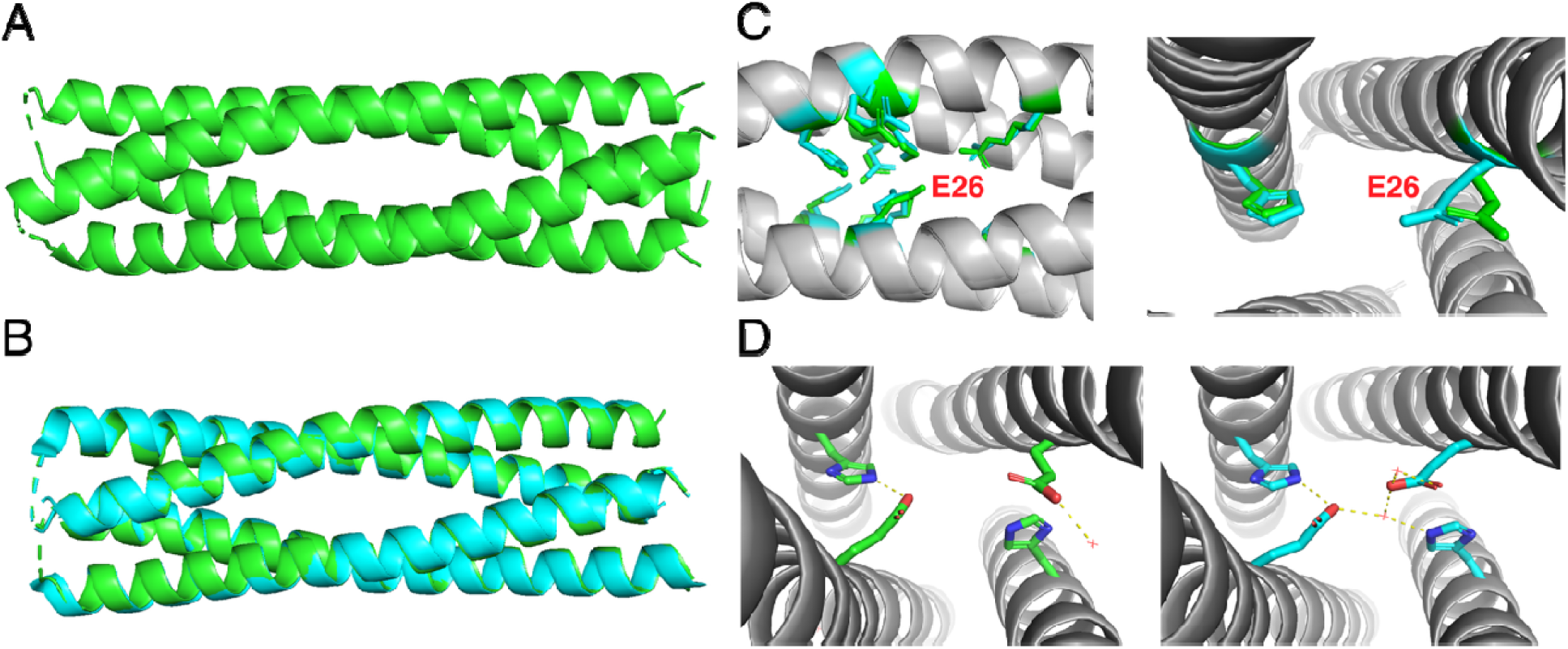
Structure of Syn-F4-Link. (A) Crystal structure of Syn-F4-Link K4Tx2. (B) Syn-F4-Link (green) and Syn-F4 (cyan) have an overall C-alpha RMSD of 0.45 Å. (C) Catalytic residues, with Syn-F4-Link residues in green and Syn-F4 residues in cyan. Glu26 residue is highlighted in red. (D) H-bond and salt bridge network among Glu26, His74 and water molecules in Syn-F4-Link (left) and unlinked Syn-F4 (right).

In addition to the crystallographic studies, we performed biophysical characterizations in solution. Analytical ultracentrifugation and size exclusion chromatography both demonstrated that Syn-F4-Link and the noncovalent dimer of Syn-F4 have similar apparent molecular weights in solution (**SI Figure 2A**). Moreover, small-angle X-ray scattering (SAXS) and 1D NMR suggest the solution structures of the covalent and non-covalent dimers are similar (**SI Figure 2B, 3, SI Table 2**). Therefore, we concluded that while biophysical studies suggest Syn-F4 and Syn-F4-Link have similar overall structures, the crystal structure of Syn-F4-Link indicates a change of conformation of the catalytic residues, which may disrupt enzymatic activity.

**Figure 3.**
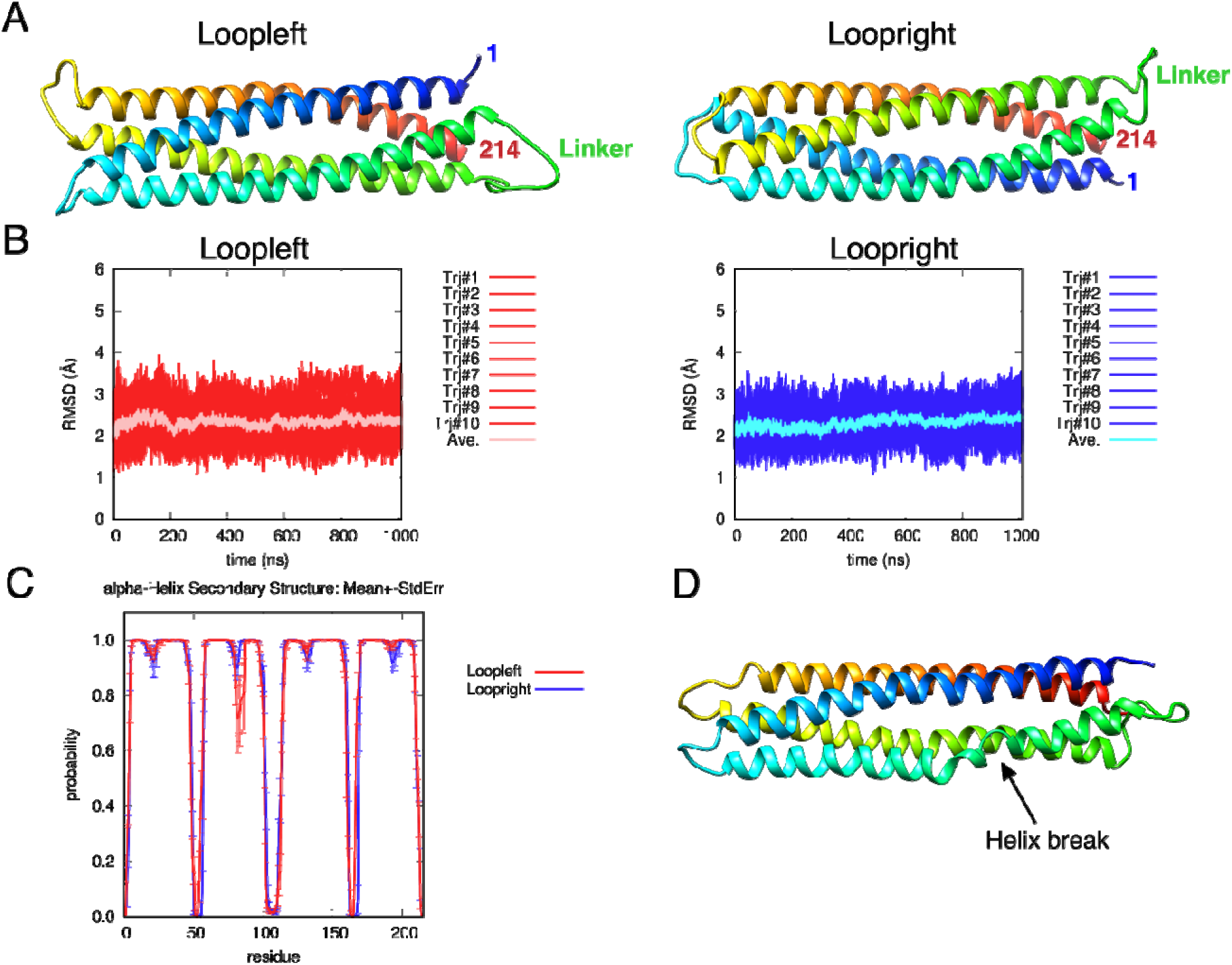
MD simulations of Syn-F4-Link. (A) Syn-F4-Link with different loop connections. N-terminus of the protein is in blue, and C-terminus is in red. Residue 1 and 214 in the chains are indicated. The covalent flexible linker is shown in green. (B) RMSD of Syn-F4-Link structures from the original models in the simulation. (C) Probability of forming an α-helical structure for each residue in MD simulations. (D) The second helix in loopleft conformation is fragile.

### Molecular dynamics (MD) simulations of Syn-F4-Link

The crystallographic data obtained for Syn-F4-Link does not reveal the structures of the loops connecting the α-helices. There is insufficient electron density, both for the turns connecting successive helices and for the residues engineered to link two monomers into a dimer. This ambiguity concerning inter-helical regions was also true for the parental protein (Syn-F4). In the case of the parental protein, we performed MD simulations, which ultimately showed that the structure with loopleft connections was more stable than the structure with the loopright connections.^26^

To compare the topology of the current structure for Syn-F4-Link with the parental homo-dimeric protein, we constructed two models for the loop connections, indicated by loopleft and loopright in **Figure 3A**. Both models were subjected to MD simulations. The results indicate that for the linked protein, loopleft and loopright have similar overall stabilities (**Figure 3B**). However, in the loopleft simulation, residues 57-101 in the second helix are fragile, allowing a break in the helix in traj09 frame101 (**Figure 3C, D**).

Furthermore, the results suggest that the linked protein with loopleft and loopright (average RMSD: ∼2 Å) are less dynamic than the unlinked parental protein with loopleft and loopright (average RMSD: ∼3 Å and ∼3.5 Å, respectively) (**SI Figure 4**). This change in dynamics may affect the active site and thereby account for the loss of catalytic activity associated with the covalent linkage of two subunits.

**Figure 4.**
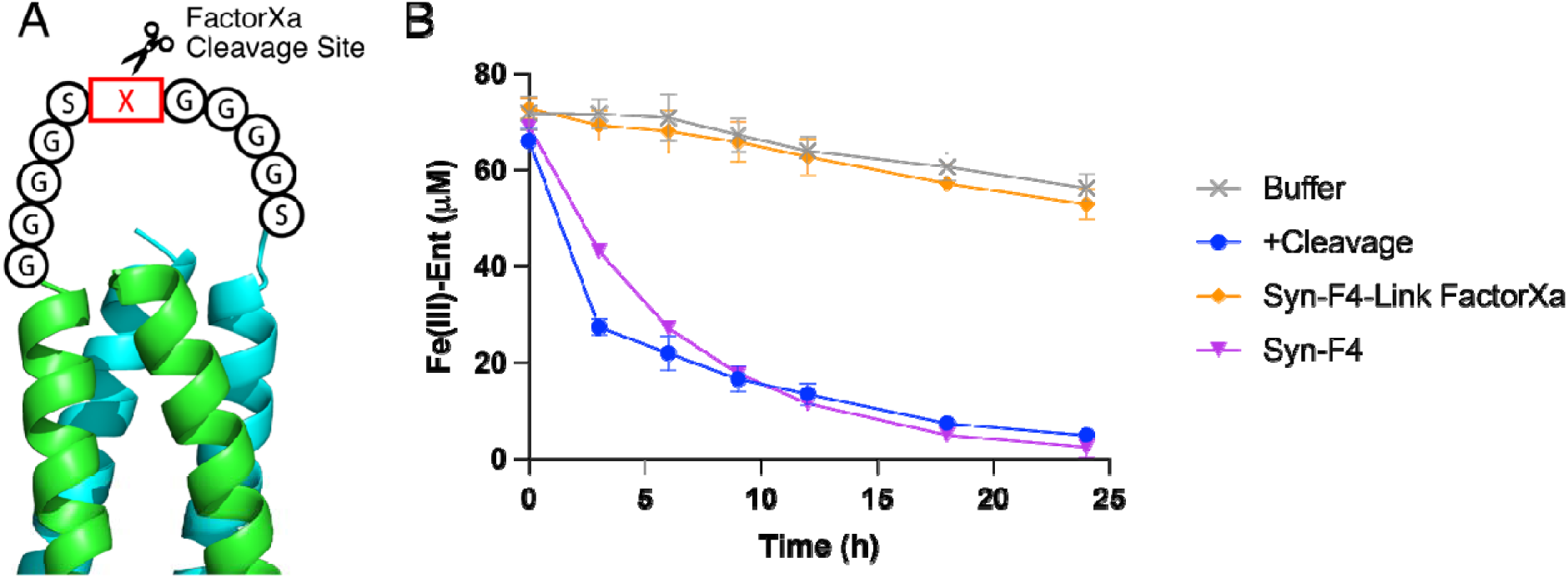
Cleavage and refolding of the linker of Syn-F4-Link derivative restores the Fe(III)-Ent hydrolysis activity. (A) Design of Syn-F4-Link with FactorXa cleavable site. (B) Hydrolysis of Fe(III)-Ent over 24 hours by Syn-F4-Link without cleavage, with cleavage and refolding, unlinked Syn-F4 and buffer.

We also probed the binding of Syn-F4-Link to Fe(III)-Ent by docking simulation. We found that while the linked protein could bind to Fe(III)-Ent in its hydrophilic pocket, the distribution of Fe^3+^ in Syn-F4-Link is distant from the catalytic residues H74 and H186 (**SI Figure 5**).

**Figure 5.**
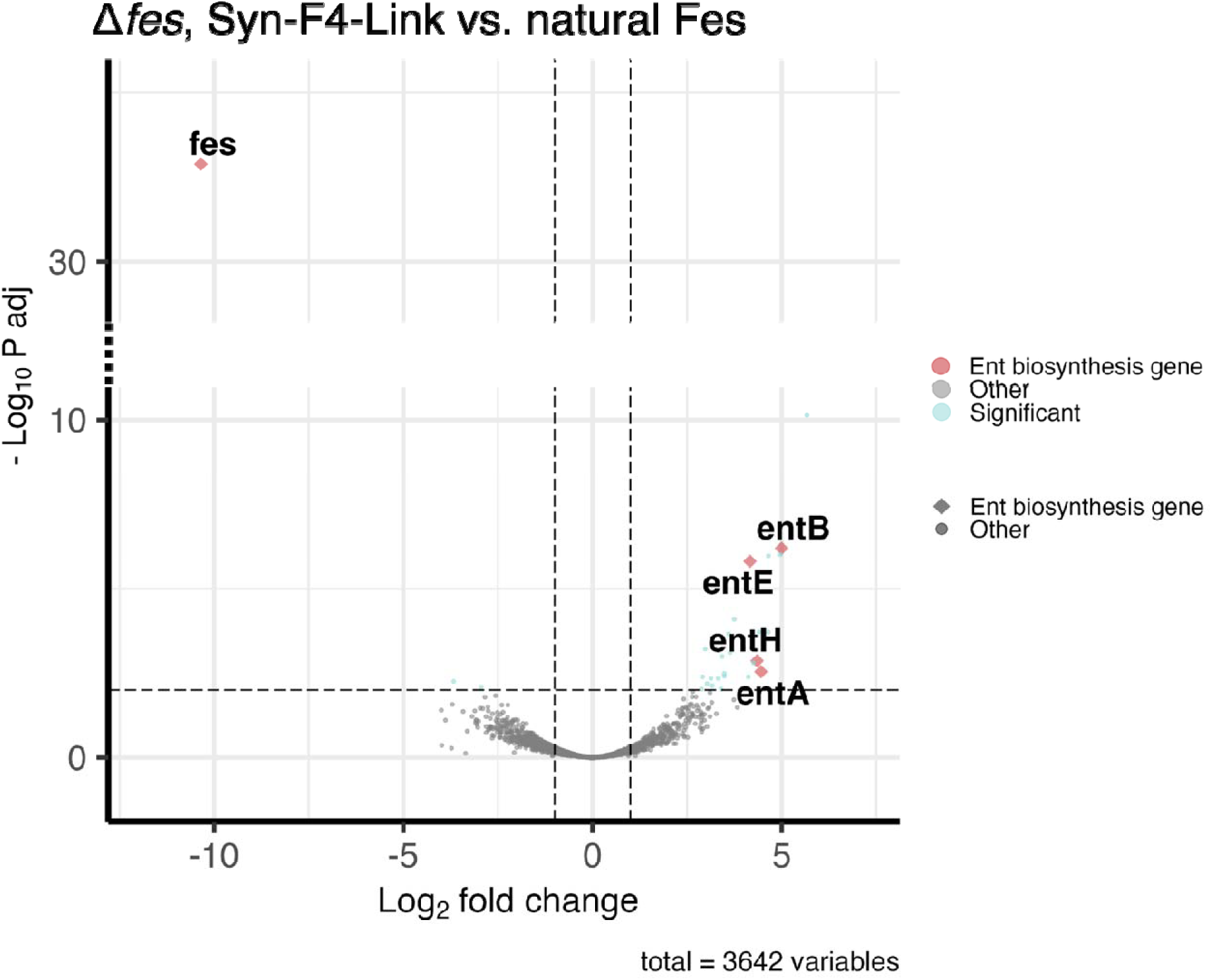
Syn-F4-Link significantly upregulates expression of enterobactin synthesis genes. The effect of Syn-F4-Link relative to natural Fes on the transcriptome of Δ*fes* while growing in minimal media is shown. Enterobactin synthesis-related genes, and *fes* gene as an internal control, are highlighted.

Together, the MD simulations suggest that Syn-F4-Link adopts a different loop connection and is less dynamic comparing to the parental Syn-F4 protein. In addition, while Syn-F4-Link still shows binding to Fe(III)-Ent, it could not position the substrate properly in the active site. These differences between Syn-F4-Link and Syn-F4 could impact the catalytic activity of the linked protein.

### Covalent linkage of two Syn-F4 chains destroys enzymatic activity

To probe the biological activity of Syn-F4-Link, we transformed a plasmid encoding Syn-F4-Link into Δ*fes* cells and attempted to grow the transformed cells in minimal media. We discovered that the Syn-F4-Link covalent dimer rescues Δ*fes* cells in iron-deficient medium, although at a slower rate than the unlinked dimer, Syn-F4 (**Figure 1B**). Next, we purified the Syn-F4-Link protein and assessed its enzymatic activity *in vitro*. Surprisingly, we found that purified Syn-F4-Link does *not* hydrolyze Fe(III)-Ent. As shown in **Figure 1C**, the parental protein Syn-F4 leads to nearly a complete loss of Fe(III)-Ent (rightmost peak in the chromatogram). However, samples containing Syn-F4-Link retain the same amount of Fe(III)-Ent as the buffer control, with breakdown products observed only at the level as seen for spontaneous hydrolysis in the control. In addition, the soluble lysate of Δ*fes* overexpressing Syn-F4-Link cannot hydrolyze Fe(III)-Ent at a rate faster than background hydrolysis (**SI Figure 6**). Thus, although the linked protein can still rescue Δ*fes* cells *in vivo*, surprisingly, it has no detectable enzymatic activity *in vitro*.

**Figure 6.**
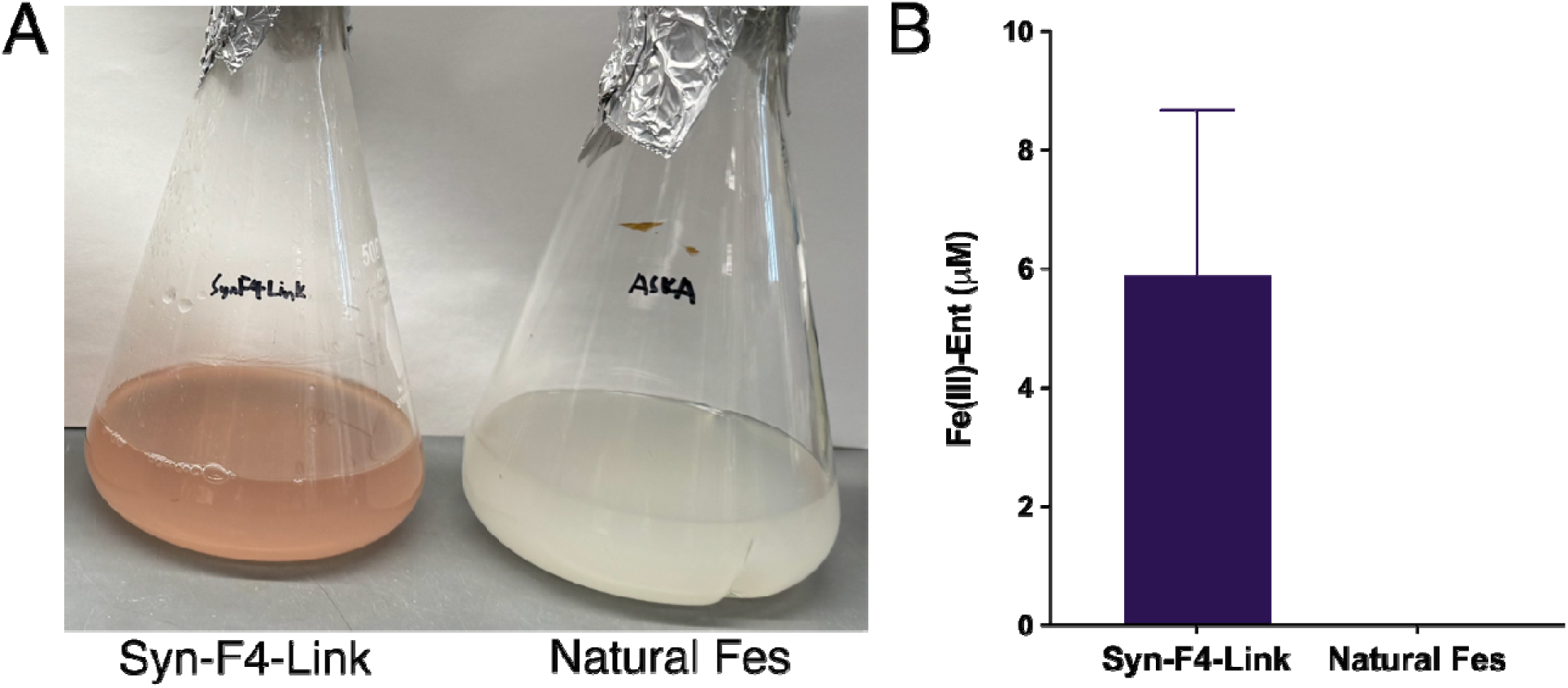
Syn-F4-Link upregulates enterobactin production. (A) Growth of Δ*fes* Syn-F4-Link and Δ*fes* natural Fes in minimal media supplied with 100 µM Fe^3+^. (B) Enterobactin abundance in media determined by reverse-phase HPLC.

### The peptide linker prevents catalytic activity of the linked protein

To further probe the surprising impact of the linker on catalytic activity, we engineered a FactorXa protease site into the linker sequence, allowing us to break the covalent linkage in the purified Syn-F4-Link protein (**Figure 4A**), and thereby enable direct comparison of enzymatic activity pre- and post-cleavage. This new variant is called Syn-F4-Link-FactorXa, and contains two copies of Syn-F4 joined by the 14-residue linker Gly-Gly-Gly-Gly-Ser-<u>Ile-Glu-Gly-Arg</u>-Gly-Gly-Gly-Gly-Ser, containing the (underlined) recognition site for the FactorXa protease.

Enzyme activity assays of this new protein showed that like the original Syn-F4-Link, purified Syn-F4-Link-FactorXa cannot hydrolyze Fe(III)-Ent (**Figure 4B**). However, after cleaving the linker with FactorXa protease, the cleaved product gained partial catalytic activity (**SI Figure 7**). We surmised that the partial (rather than complete) return of activity might indicate that although the linker was cut, most of the dimeric protein was kinetically trapped in the conformation it had assumed prior to proteolytic cleavage. To test this possibility, we disassembled and denatured the cleaved products by running the material over reverse phase HPLC and eluting with organic solvents. We then lyophilized the sample to remove acetonitrile and refolded the protein in a non-denaturing buffer (75mM HEPES, 1M NaCl, pH 8.0). We found that refolding the cleaved products fully restored the enzymatic activity (**Figure 4B**). These results demonstrate that covalently linking two chains of Syn-F4 – either with the original decapeptide linker or with the modified 14 residue peptide linker with a FactorXa cleavage site – destroys the catalytic activity of the *de novo* protein.

**Figure 7.**
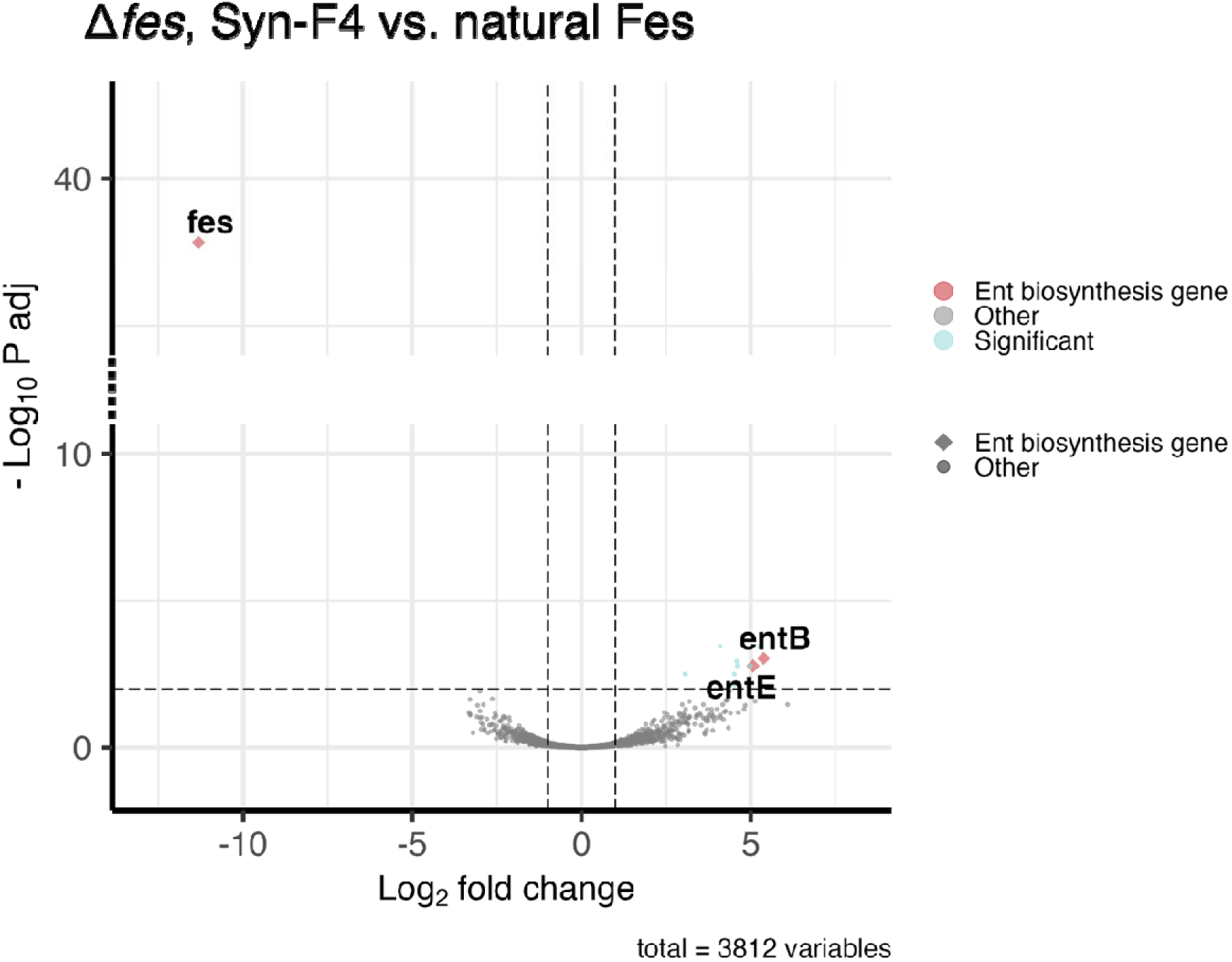
Syn-F4 significantly upregulates expression of enterobactin synthesis genes. The effect of Syn-F4 relative to natural Fes on the transcriptome of Δ*fes* in minimal media is shown. *entB* and *entE* genes, and the internal control gene *fes*, are highlighted.

### The linked protein upregulates enterobactin biosynthesis in *E. coli*

The parental Syn-F4 protein rescues Δ*fes* cells *in vivo,* and *in vitro* it catalyzes the same reaction as the natural Fes enzyme. On the other hand, Syn-F4-Link rescues Δ*fes* cells *in vivo,* but *fails* to catalyze the hydrolysis of Fe(III)-Ent. Therefore, the linked protein must enable the growth of Δ*fes* cells by some alternative – perhaps regulatory – mechanism. To probe this possibility, we performed global transcriptomic analysis of *E. coli* cells expressing Syn-F4 or Syn-F4-Link and compared these results to cells expressing the natural Fes protein. RNA-seq analysis of Δ*fes* cells expressing Syn-F4-Link revealed that expression of the covalently linked dimer upregulates enterobactin biosynthesis genes, including *entA*, *entB*, *entE*, *entF*, and *entH* (**Figure 5**). Importantly, this upregulation is also observed in a pseudo-wildtype strain (Keio parent strain, rather than Keio Δ*fes*) expressing the natural Fes protein. This last result demonstrates that expression of Syn-F4-Link – not just background starvation for iron – causes the observed upregulation of genes related to enterobactin biosynthesis and iron assimilation (**SI Figure 8**).

These transcriptomic results indicate that Syn-F4-Link leads to overproduction of enterobactin. Therefore, expression of Syn-F4-Link in Δ*fes* cells, which cannot degrade the siderophore, should lead to an accumulation of Fe(III)-Ent. To test this expectation, we grew cultures of Δ*fes* cells overexpressing Syn-F4-Link in iron-enriched media and compared them to the same cells harboring the natural *fes* gene. As shown in **Figure 6A**, Δ*fes* cells expressing Syn-F4-Link produce red cultures, consistent with the accumulation of Fe(III)-Ent. Moreover, quantification of the siderophore by HPLC showed that Syn-F4-Link induces the synthesis of substantially more Fe(III)-Ent than does the natural Fes protein (**Figure 6B)**.

### The parental homodimeric *de novo* protein Syn-F4 has two biological functions

After discovering that Syn-F4-Link is not enzymatically active, but instead rescues Δ*fes* cells by a regulatory mechanism that induces overproduction of enterobactin, we questioned whether the enzymatically active parental protein, Syn-F4, might also have a secondary activity upregulating the biosynthesis of enterobactin. To address this question, we analyzed the effect of Syn-F4 expression on the transcriptome. The results, shown in **Figure 7 and SI Figure 9**, indicate that Syn-F4 does indeed induce increased expression of enterobactin biosynthesis genes. Thus, the original Syn-F4 homodimer is a bifunctional *de novo* protein. It can rescue Δ*fes* by performing the deleted enzymatic function to hydrolyze Fe(III)-Ent, and it also alters gene expression and upregulates enterobactin biosynthesis.

## DISCUSSION

Several years ago, we described Syn-F4 as the first *de novo* protein that shares no ancestry with natural enzymes, but nonetheless provides a life-sustaining catalytic function *in vitro* and *in vivo*.^25^ Subsequent determination of the crystal structure of Syn-F4 revealed a 4-helix bundle comprising two α-helical hairpins arranged in a *syn* (as compared to *anti*) topology.^26^ Because this orientation placed the C-terminus of one chain close to the N-terminus of the second chain, we reasoned that linking these termini would enhance (or at least not destroy) activity by creating a single chain that could fold into its active conformation without paying the entropic price of dimerization.

As described above, the linked protein was constructed and shown to fold into a structure that closely resembled the original homodimer. Moreover, as expected, it rescued the growth of Δ*fes* cells in minimal media. Surprisingly, however, the purified Syn-F4-Link protein was *not* enzymatically active and failed to hydrolyze ferric enterobactin. Instead, we found that Syn-F4-Link rescues Δ*fes* cells by using a regulatory, rather than enzymatic, mechanism.

This surprising change in the mechanism of biological rescue led to a closer examination of the structure and dynamics of Syn-F4-Link relative to its progenitor Syn-F4. Specifically, MD simulations of Syn-F4-Link with either of two possible loop connections (loopleft or loopright) showed that a loopleft connection like that in the unlinked Syn-F4 homodimer would cause the second helix to be unstable. This finding suggests that the engineered covalent linkage favors folding into the inactive loopright conformation. In this loopright conformation, the hydrogen-bonding network among the active site residues is disrupted, which would account for the loss of catalytic activity. In addition, the MD simulation indicated Syn-F4-Link was less dynamic than the unlinked Syn-F4, which would hinder the ability of the linked chain from sampling the catalytically active confirmation. This finding suggests that for this *de novo* enzyme, as for many natural enzymes, structural dynamics are important for catalytic activity.^27,28^

It was surprising that linking the two chains abrogated enzymatic activity. It would be even more surprising if this linkage simultaneously generated an entirely new regulatory activity that was not present in the parental homodimer. Therefore, we questioned whether the Syn-F4 homodimer might also upregulate enterobactin biosynthesis. As shown in **Figure 7**, this turned out to be true: The parental Syn-F4 homodimer – like the engineered Syn-F4-Link protein – upregulates enterobactin biosynthesis. This regulatory function of Syn-F4 was not observed previously because it is latent when the *de novo* rescuer acts as an enzyme, but was revealed when enzymatic activity was destroyed by engineering a covalent linkage joining the two chains.

Our initial isolation of numerous *de novo* proteins that rescue Δ*fes* cells,^18^ followed by subsequent genetic, biochemical and structural characterization of the linked and unlinked versions of Syn-F4 reveal that it is not difficult to find non-natural proteins that enable the growth of Δ*fes* cells in iron-limited conditions. Importantly, these studies reveal that novel proteins that did not arise in nature can use either (or both) catalytic or regulatory mechanisms to sustain the growth of living organisms.

While Syn-F4-Link significantly upregulates enterobactin biosynthesis, this alone is not sufficient to explain how the cell acquires sufficient Fe^3+^ to survive in iron deficient media. We hypothesize that binding of Syn-F4-Link to Fe(III)-Ent could disrupt the precise coordination between enterobactin and Fe^3+^, thereby releasing iron for cellular metabolism. Likely, this process is inefficient. Therefore, only when Fe(III)-Ent is overexpressed could the cell gain enough iron for survival, which explains the importance of Syn-F4-Link in upregulating the biosynthesis of Fe(III)-Ent. Future studies characterizing the binding of enterobactin to iron under the effect of Syn-F4-Link could enhance our understanding of the *de novo* protein’s life-sustaining mechanism.

## ACKNOWLEDGEMENTS

We thank J. Eng for assistance with mass spectrometry and I. Pelczer for assistance with protein NMR. We thank all members of the Hecht lab for helpful discussion. This work was funded by NSF MCB-1947720.

## AUTHOR CONTRIBUTIONS

G.L., S.T. and, M.H.H. initiated the project idea. S.T. constructed Syn-F4-Link. G.L. performed *in vivo* rescue, *in vitro* protein biophysics and enzymology, RNA-seq, and metabolomics assay. M.N., K.K., and R.A. conducted crystallography and SAXS. K.U. conducted MD and docking simulations. G.L. and M.H.H. drafted the manuscript, with input from all the authors.

## MATERIALS AND METHODS

### Gene construction

To generate the Syn-F4-link construct, in which two Syn-F4 coding sequences are joined by a flexible linker (GGGGSGGGGS), two overlapping DNA fragments were amplified by PCR using the Syn-F4-containing plasmid (Syn-F4 in p3glar) as the template. The first fragment was amplified with primers F-1 and F-2, introducing the linker sequence at the 5′ end of the Syn-F4 coding region. The second fragment was amplified using primers V-1 and V-2; this removed the stop codon of Syn-F4 and appended the same linker sequence to its 3′ end. Additionally, the second fragment included vector-derived sequences flanking the Syn-F4-linker region, allowing it to serve as the backbone for plasmid assembly. The two fragments were assembled using Gibson Assembly (NEB), transformed into *E. coli* DH5α, and verified by colony PCR and Sanger sequencing.

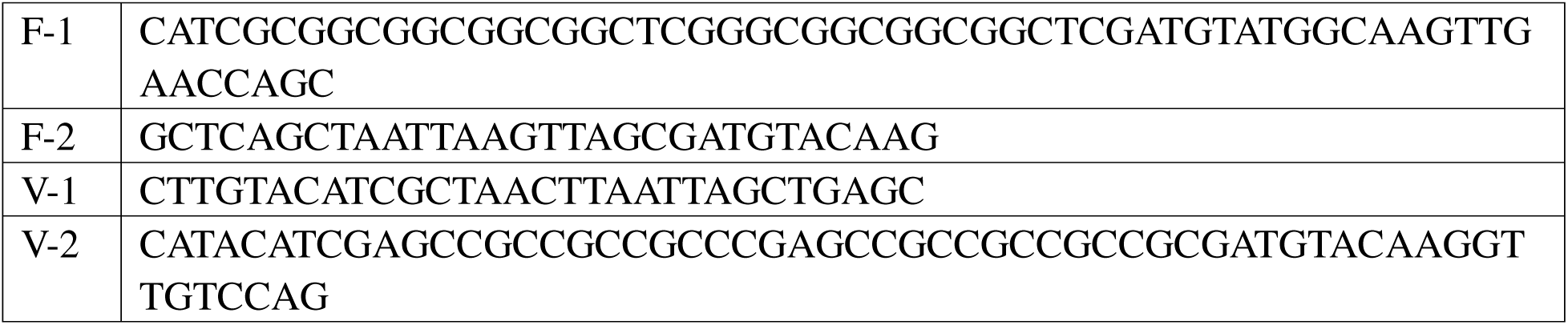

### Protein expression and purification

Syn-F4 (K4T), Syn-F4-Link (K4Tx2) and Syn-F4-Link (FactorXa) genes were cloned into the expression vector p3glar, with chloramphenicol resistance. Proteins were overexpressed in BL21 (New England Biolabs), where expression was induced with 0.1 mM IPTG at OD_600_ ∼0.5 and incubated at 18 °C for 18 hours. Cells were harvested by centrifugation, resuspended in purification buffer (75 mM HEPES, 1 M NaCl, pH 8.0), and lysed by sonication (10 seconds on, 50 seconds off, total sonication time of 3 minutes 30 seconds). The soluble lysate was separated from the insoluble fraction by centrifugation and filtered through a 0.45 μm syringe filter (Millipore). The soluble lysate containing overexpressed protein was loaded onto a nickel column (HisTrap HP, Cytiva), washed with purification buffer containing 50 mM imidazole, and eluted with 375 mM imidazole. The eluted fraction was then loaded on a size exclusion chromatographic column (HiLoad 26/600 Superdex 75 pg, Cytiva) and fractions corresponding to the expected molecular weight of the protein were combined and stored at −80 °C. The purity of protein was confirmed by SDS-PAGE.

### Crystallography and structure determination

For crystallization, the Syn-F4-Link (K4Tx2) protein was purified by immobilized metal ion affinity chromatography with COSMOGEL His-Accept (Nacalai Tesque) (equilibration/wash buffer: 50 mM sodium phosphate buffer (pH 8.0) containing 300 mM NaCl and 10% glycerol; elution buffer: 50 mM sodium phosphate buffer (pH 8.0) containing 300 mM NaCl, 10% glycerol and 250 mM imidazole). The protein was further purified by anion exchange chromatography (25 mM Tris-HCl buffer (pH 8.5) containing 10% glycerol with a linear gradient of NaCl from 0 to 1.5 M) with a RESOURCE Q 6 mL column (Cytiva) and size-exclusion chromatography (25 mM Tris-HCl buffer (pH 9.0) containing 300 mM NaCl, 10% glycerol and 200 mM Arg-HCl with a Superdex 75 Increase 10/300 GL column (Cytiva).

Plate-like crystals were obtained in a drop composed of 1 μL of the protein solution (9.2 mg/mL) and 1 μL of the reservoir solution (110 mM sodium malonate, pH 7.0, 9.5% w/v polyethylene glycol 3,350) by the hanging drop vapor diffusion method against 500 μL of the reservoir solution with a 24-well plate at 20°C in a week.

X-ray diffraction data were collected at the Photon Factory, BL-17A (KEK, Tsukuba, Japan) with a EIGER X 16M detector (Dectris, Baden, Switzerland). The data collection was carried out at 95 K with 30% glycerol as a cryoprotectant. Diffraction data were processed with the programs XDS^29,30^ and AIMLESS^31^. The structure was solved by molecular replacement method using Phaser with a model structure of Syn-F4 (K4T) (PDB ID: 8H7D)^26^. The model was corrected with the program COOT^32^ and was refined with the program REFMAC5^33^ in the CCP4 suite^34^. The quality of the model was inspected by the programs PROCHECK^35^ and MolProbity^36^. All statistics are presented in **SI Table 1**. The atomic coordinates and the structure factors have been deposited in the Protein Data Bank with the accession code 9UGR.

### Small-angle X-ray scattering (SAXS)

SAXS measurements were performed for samples (∼5 mg/mL) of Syn-F4-Link (K4Tx2) and Syn-F4 (K4T) and hen egg lysozyme dissolved in 25 mM MES buffer (pH 6.5) containing 100 mM NaCl, 10% glycerol and 200 mM Arg-HCl at 20 °C using synchrotron radiation (λ = 1.3 Å) at the Photon Factory BL-10C beamline^37^ (KEK, Tsukuba, Japan) with a PILATUS3 2M detector (Dectris, Baden, Switzerland) at a sample-detector distance of 1 m. The two-dimensional scattering images were integrated into one-dimensional scattering intensities *I*(*q*) as a function of the magnitude of the scattering vector *q* = (4π/λ)sin(θ/2) using SAngler^38^, where θ is the total scattering angle. The forward scattering intensity, *I*(*q*→0), and radius of gyration, *R*_g_, were estimated by Guinier approximation ^39^ using AUTORG in ATSAS^40^ with SAngler^38^. Forward scattering intensity normalised by the protein concentration (mg/mL), *I*(*q*→0)/*c*, is proportional to weight average molecular mass (*M*_w_). Lysozyme (*M*_w_ = 14.3 kDa) was used as a reference standard of the molecular mass. A low-resolution dummy atom model was constructed from the SAXS data using *ab initio* shape modelling programs in the ATSAS program suite^40^. Calculations of rapid *ab initio* shape determination were performed ten times by DAMMIF^41^ without a symmetry constraint, and the generated models were aligned and averaged by DAMAVER^42^. The averaged model was modified with the fixed core by DAMSTART and further refinement of the model was performed by DAMMIN^43^. Superimposing the dummy atom model and crystal structures was performed by SUPCOMB^44^. The SAXS data and dummy-atom models of Syn-F4-Link (K4Tx2) and Syn-F4 (K4T) have been deposited in Small Angle Scattering Biological Data Bank^45^ (SASBDB accession codes: xxxxxxx, yyyyyyyy).

### Molecular dynamics and docking simulations

All-atom MD simulations were performed for the two Syn-F4-Link constructs with different loop connections (loop-left and loop-right) in a rectangular periodic box. The initial conformations of the loop regions were generated by MODELLAR^46^. The force field of the protein was taken from Amber ff14SB^47^, and the TIP3P water model^48^ was used. Potassium and chloride ions were added to neutralize the system and achieve a salt concentration of 100 mM. Following energy minimization, the system was heated linearly from 1 K to 310 K for 1 ns. The initial velocities of atoms were randomly assigned according to the Maxwell-Boltzmann distribution. The system was then equilibrated for 10 ns under NPT conditions (1 bar, 310 K). Production run was conducted for 1.0 μs. Snapshots were saved every 100 ps, yielding 10,000 frames per simulation. To enhance sampling, this procedure was repeated ten times with different initial velocities, generating a total of ten independent trajectories. Finally, 100,000 snapshots were collected for subsequent analyses, including secondary structure population and root-mean-square deviation (RMSD) from the initial structure. A subset of 1,000 snapshots, extracted every 10 ns, was used for the subsequent docking simulations. All simulations and analyses were carried out using the Amber18 and AmberTools18 software packages^49^.

### Enzymatic assay for hydrolysis of Fe(III)-Ent

Fe(III)-Ent was prepared as described by Donnelly *et al.*^25^ To assay catalytic activity, 25 μM of purified protein and 120 μM Fe(III)-Ent were incubated in 75 mM HEPES, 1 M NaCl, pH 8.0 at 37 °C. Aliquots were collected at various time points, quenched by 0.5 volume of 2.5 M HCl in methanol, and Fe(III)-Ent and its hydrolysis products were extracted by 2 volumes of ethyl acetate. The organic layer was analyzed by reverse-phase HPLC, and the enterobactin and hydrolysis products were monitored by absorbance at 316 nm.

### RNA-seq

Δ*fes* or Keio parent *E. coli* cells from the Keio collection^50^ harboring Syn-F4, Syn-F4-Link, natural *fes* from the ASKA collection^51^ or empty vector were incubated in M9-glucose minimal media at 37 °C. Cells were collected at OD_600_ 0.2-0.5 and resuspended in 0.5 mL DEPC treated water (Invitrogen) and kept frozen. The total RNA from cells were released by cryomill and purified by Direct-zol RNA Miniprep kits (Zymo Research). cDNA libraries were prepared by ribo-depletion followed by RNA-Seq Directional Library Prep on Apollo 324 Robot. Next-generation sequencing of the library was done with NovaSeq SP 100nt Lane v1.5 (Princeton University). The sequencing data was processed on Galaxy.^52^ The quality of RNA-seq data were tested by FastQC. The data were then processed by Trimmomatic to trim the adapters^53^, mapped to the *E. coli* K12 strain BW25113 reference genome^54^ using BWA-MEM^55^, and the genes counts were quantified by htseq-count^56^. Differential expression analysis was conducted by DESeq2^57^ with R language. The volcano plots were created by EnhancedVolcano package^58^ on R.

### Quantification of Fe(III)-Ent in culture

Δ*fes* cells expressing either Syn-F4-Link, Syn-F4, natural *fes* from the ASKA collection or S824 (a *de novo* 4-helix bundle protein with no biological function)^59^ were incubated in M9-glucose minimal media supplemented with 100 μM FeCl_3_ at 37 °C. After turbidity was observed, the cells and liquid culture were separated by centrifugation. Liquid culture was acidified by hydrochloric acid, and enterobactin was extracted by ethyl acetate. The organic layers were analyzed by reverse-phase HPLC to quantify the amount of enterobactin. The elution peak corresponding to enterobactin was collected and analyzed on mass spectrometry to confirm the identity of the molecule.

## Supplementary Information

**SI Figure 1.**
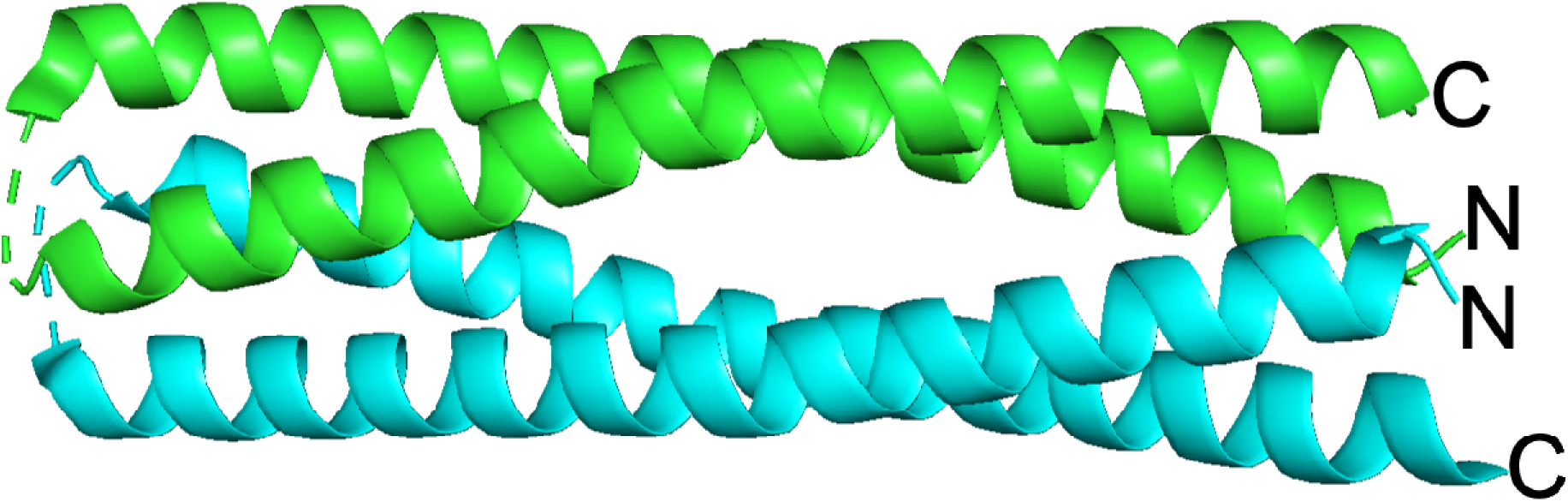
Crystal structure of Syn-F4 K4T (PDB 8H7D). N- and C-termini of each chain (green and cyan) are labeled.

**SI Figure 2.**
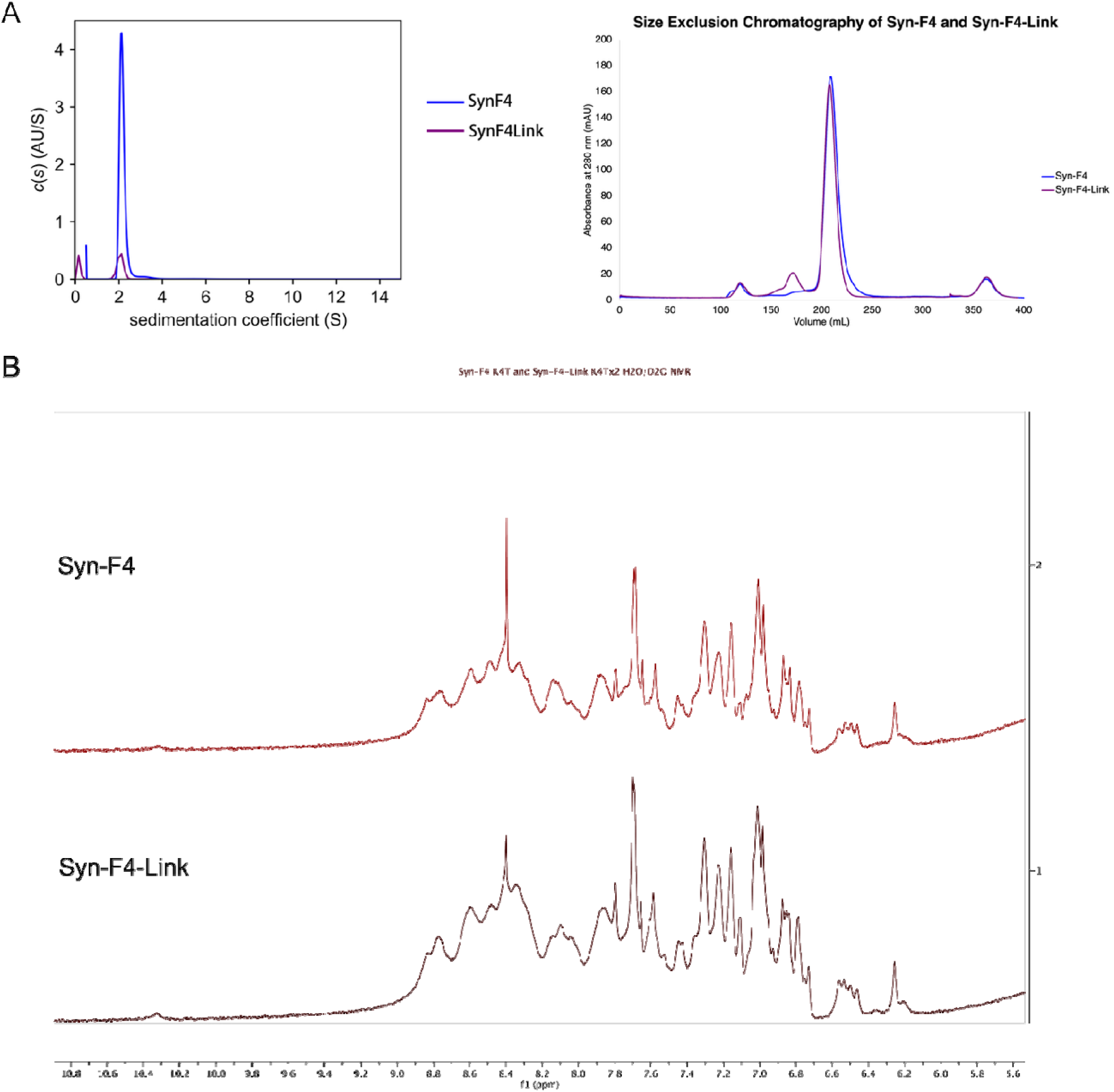
Biophysical characterization of Syn-F4-Link. (A) Analytical ultracentrifuge (left) and size exclusion chromatography (right) suggest Syn-F4-Link and Syn-F4 has similar apparent molecular weight in solution. (B) 1D NMR of Syn-F4 and Syn-F4-Link. The amide proton and tryptophan peaks are shown.

**SI Figure 3.**
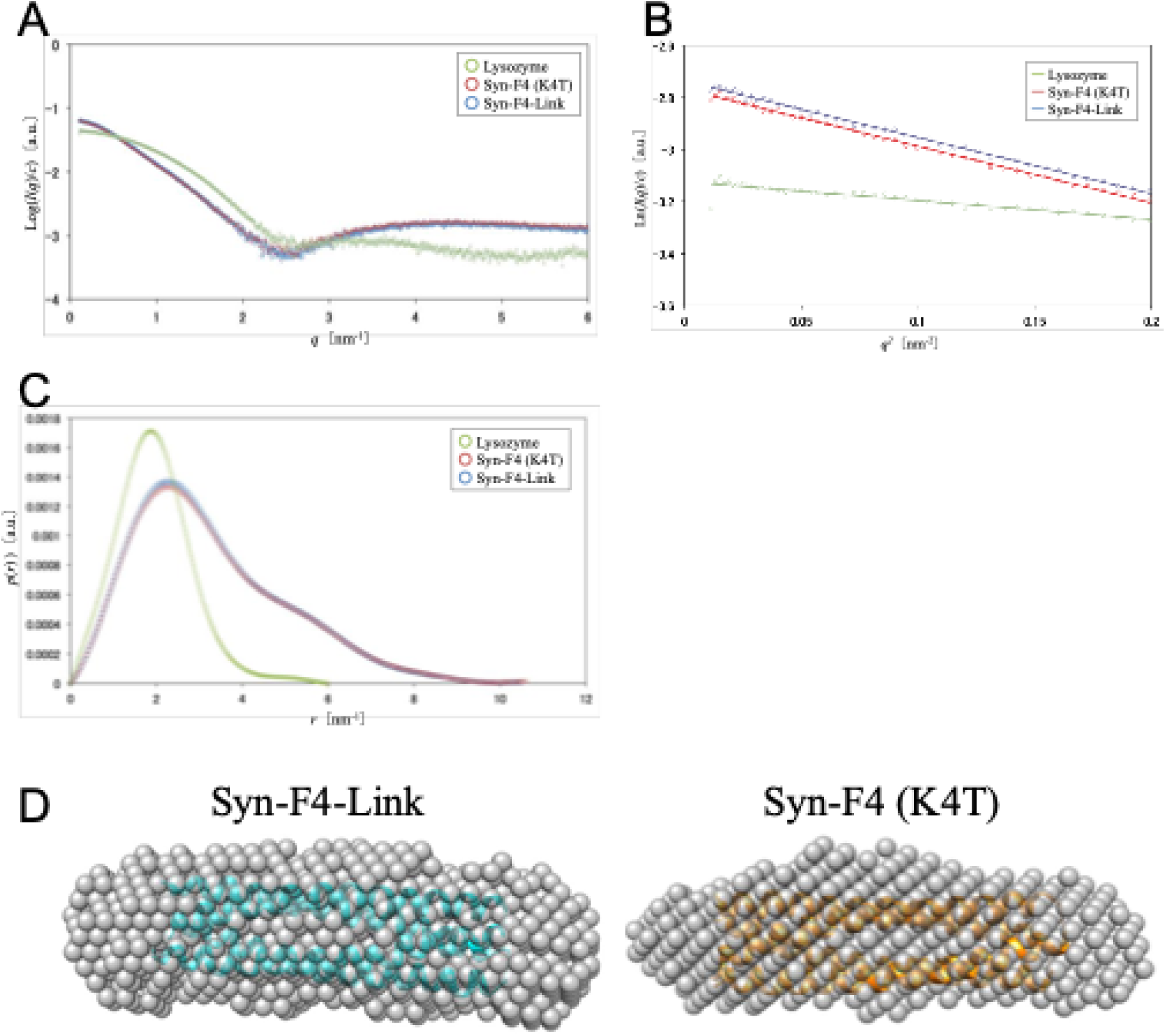
Small-angle X-ray scattering (SAXS) analyses of Syn-F4-Link and Syn-F4 (K4T). (A) Concentration-normalized scattering intensities. (B) Guinier plots. (C) Pair-distance distribution functions. Lysozyme was used as a molecular mass reference standard. SAXS analysis result and estimated molecular mass are summarized in SI Table 2. (C) Dummy atom models of Syn-F4-Link and Syn-F4 (K4T) calculated from the SAXS data using *ab initio* modeling programs, DAMMIF and DAMMIN. For reference, the crystal structures (ribbon representation) are superimposed on the dummy atom models.

**SI Figure 4.**
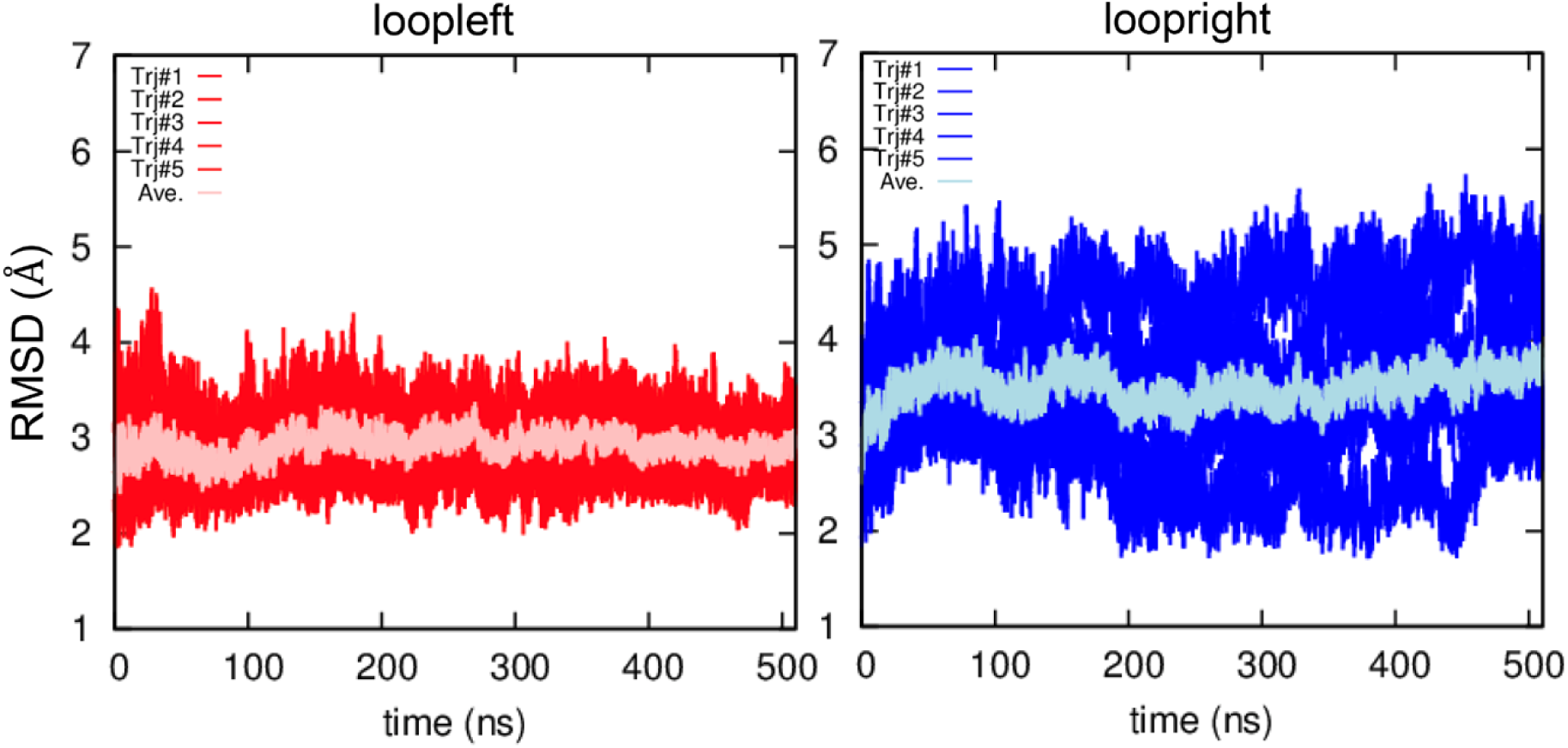
MD simulation of Syn-F4 in loopleft and loopleft conformations. Figure adopted from Kurihara et al.

**SI Figure 5.**
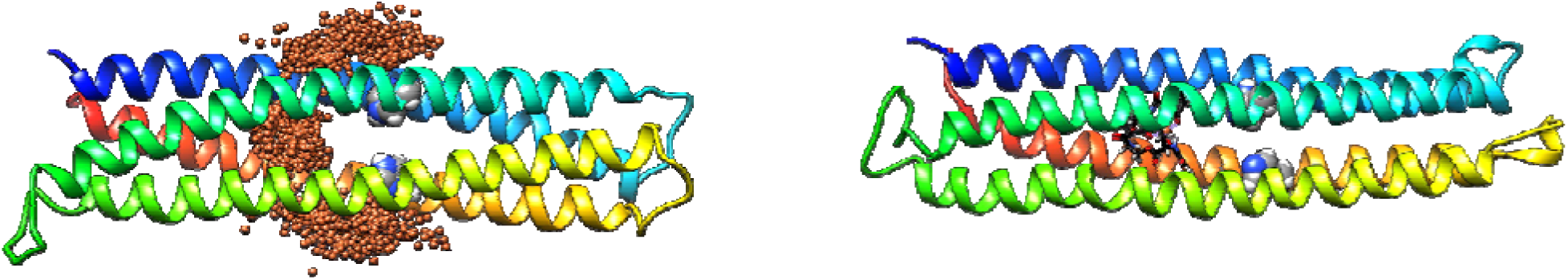
Docking simulation of Fe(III)-Ent onto Syn-F4-Link with loopright conformation. Left: distribution of Fe^3+^ in docking simulations (orange spheres). Right: the docking pose with the lowest docking score. Fe(III)-Ent molecule is shown in ribbon model. H74 and H186 are shown in blue and grey spheres.

**SI Figure 6.**
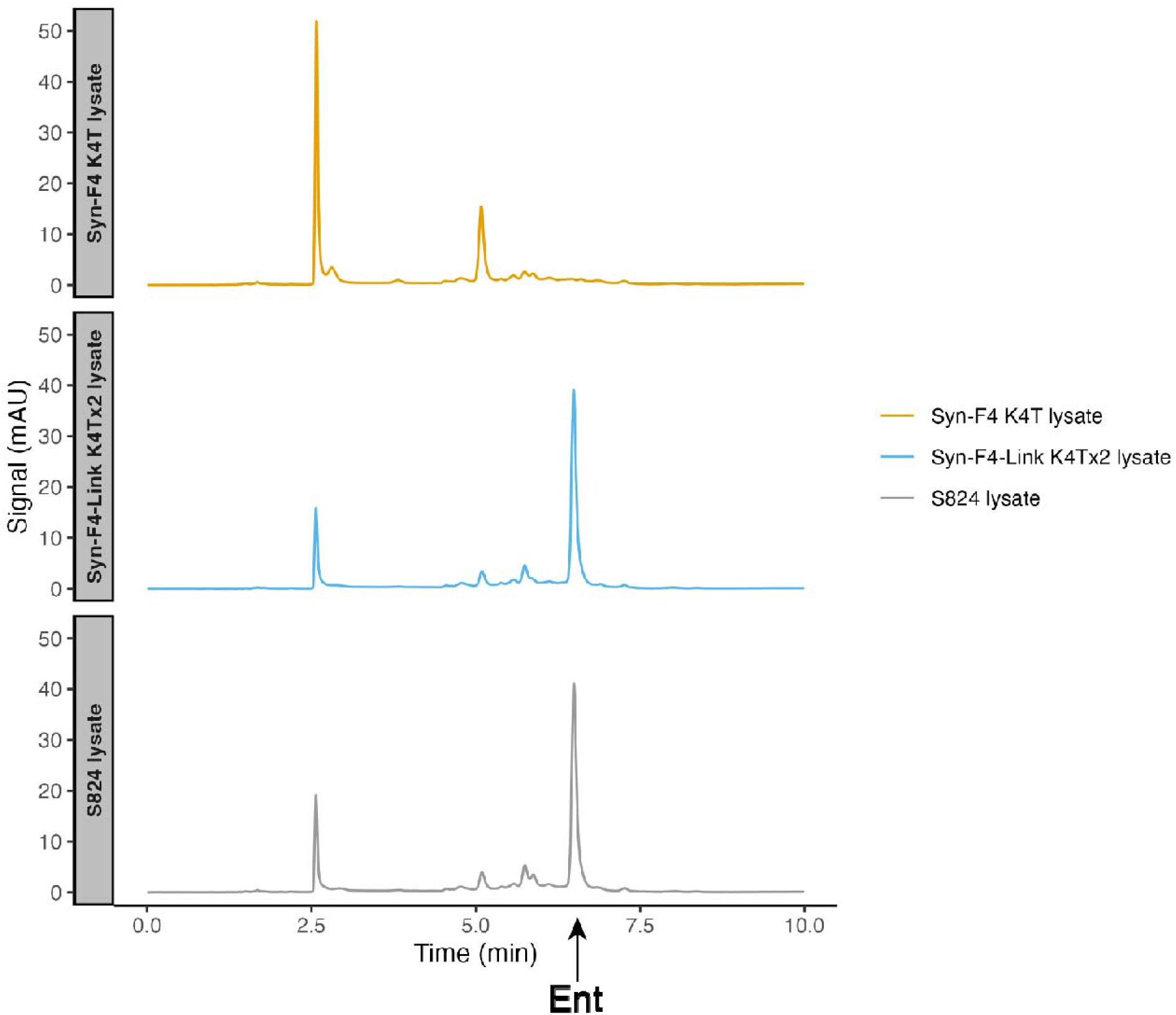
HPLC chromatogram of enterobactin and its hydrolysis products by the soluble lysate of Δ*fes* cells expressing Syn-F4, Syn-F4-Link and S824 (negative control). Peak corresponding to enterobactin (Ent) is labeled.

**SI Figure 7.**
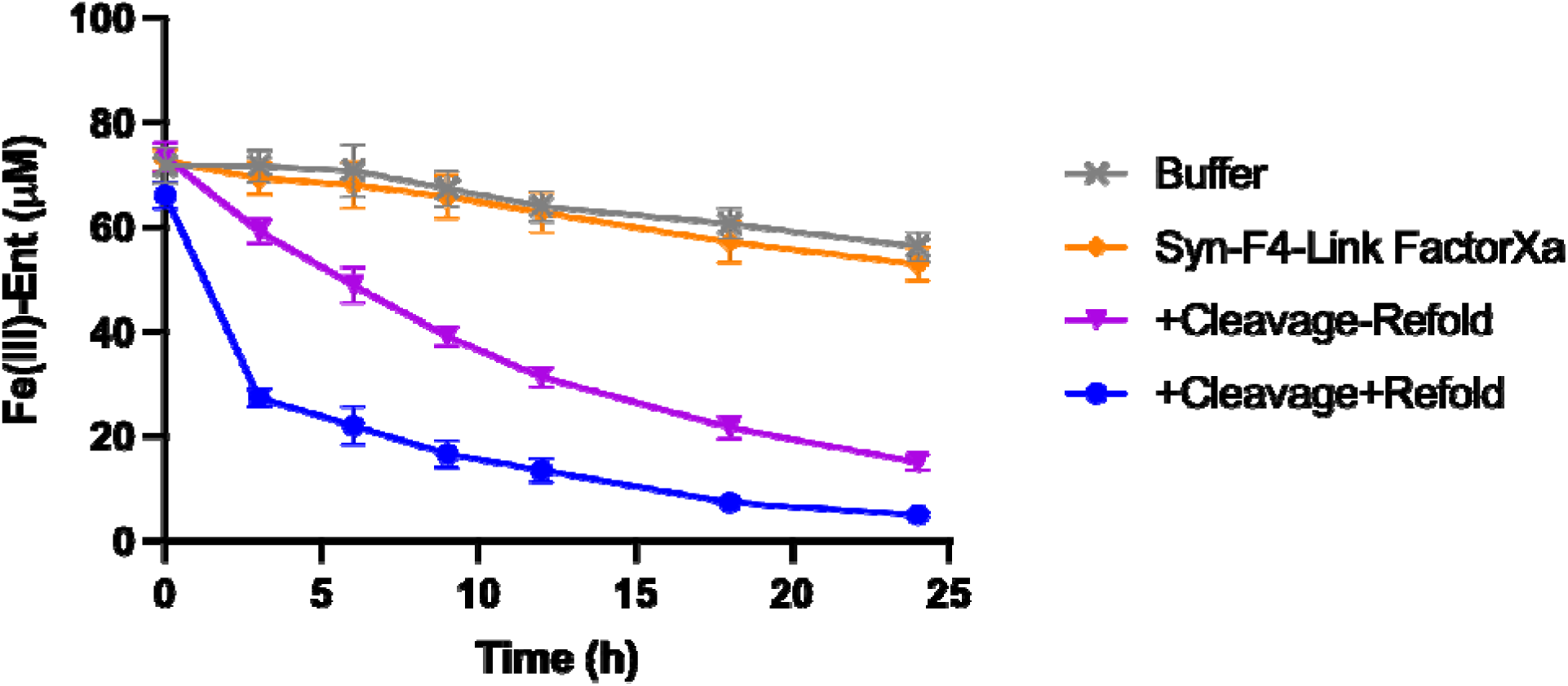
Hydrolysis of Fe(III)-Ent over 24 hours by modified Syn-F4-Link without cleavage (Syn-F4-Link FactorXa), with cleavage only (+Cleavage-Refold), with cleavage and refolding (+Cleavage+Refold) or buffer.

**SI Figure 8.**
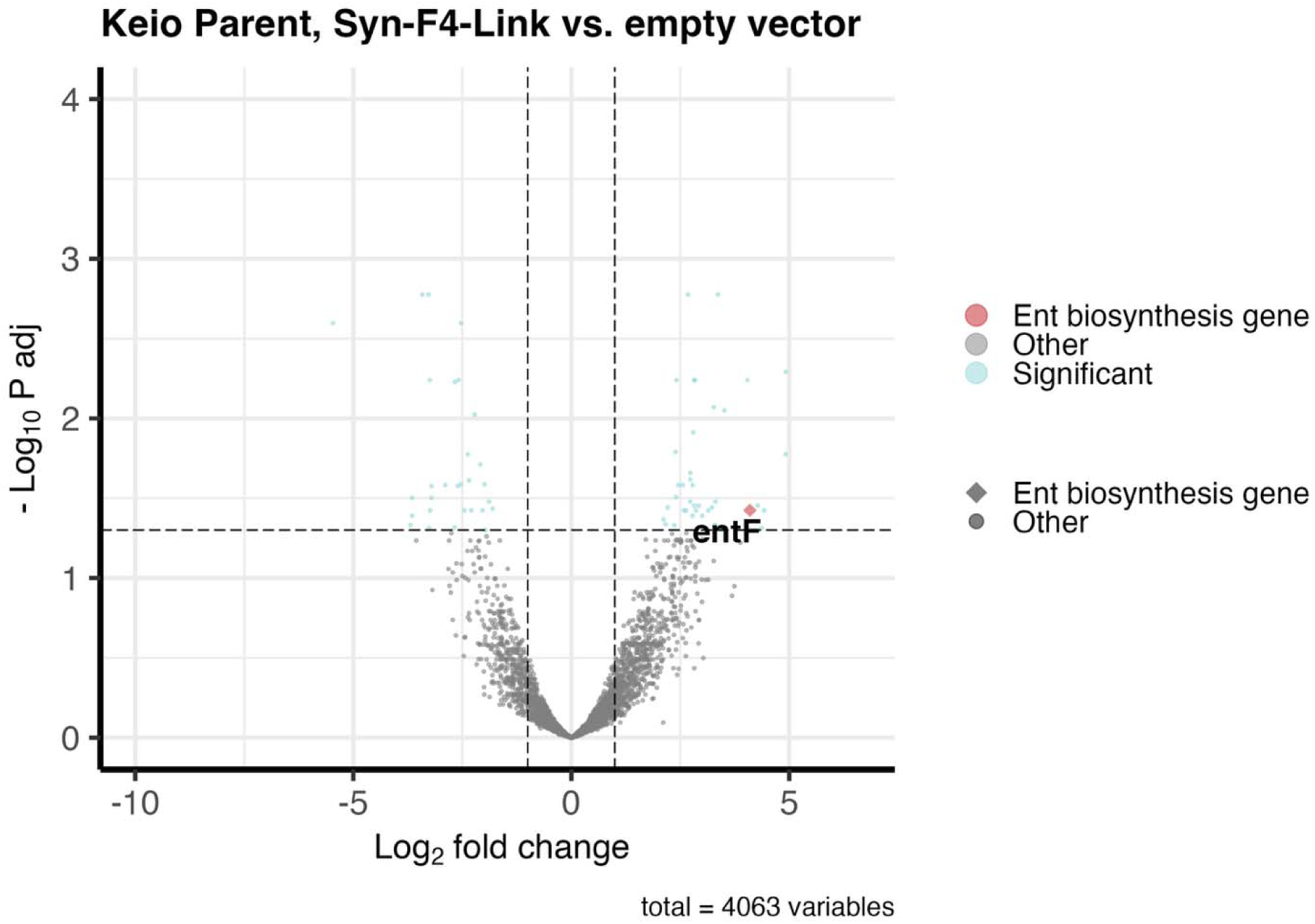
RNA-seq of Syn-F4-Link vs. empty vector in Keio parent strain. The expression of *entF* gene is significantly upregulated.

**SI Figure 9.**
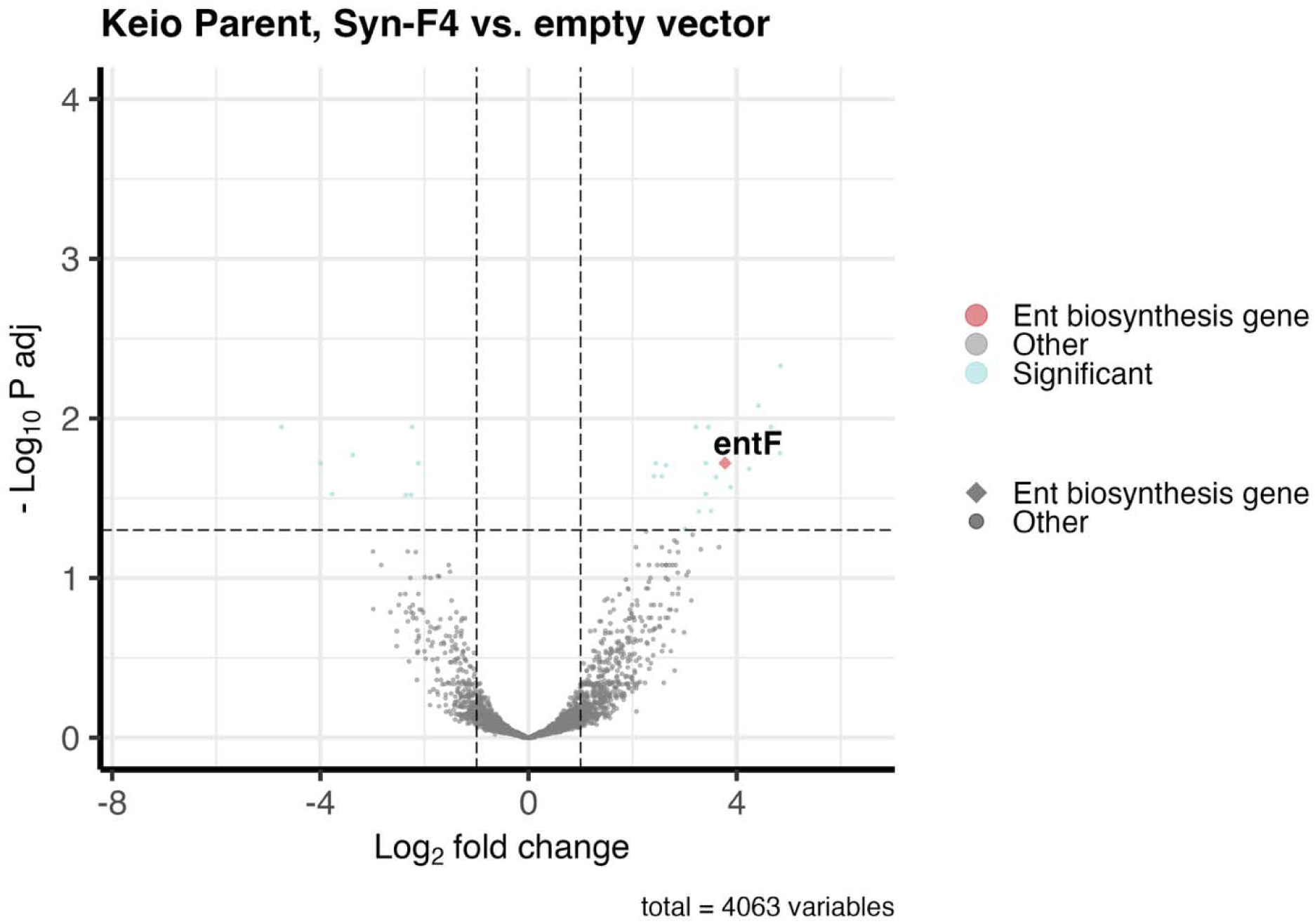
RNA-seq of Syn-F4 vs. empty vector in Keio parent strain. The expression of *entF* gene is significantly upregulated.

**SI Table 1.** X-ray data collection and refinement statistics.

| Crystal | Syn-F4-Link (K4Tx2) |
| --- | --- |
| *Data collection* |  |
| Space group | *C*2 |
| Unit-cell parameters | $a = 61.44 \text{ \AA}$, $b = 47.39 \text{ \AA}$, $c = 70.13 \text{ \AA}$<br>$\alpha = 90.00^\circ$, $\beta = 103.63^\circ$, $\gamma = 90.00^\circ$ |
| X-ray source | KEK Photon Factory (PF) BL-17A |
| Wavelength ($\text{\AA}$) | 0.980 |
| Resolution ($\text{\AA}$) | 37.12–2.30 (2.38–2.30) |
| Unique reflections | 8823 (866) |
| Average redundancy | 6.4 (6.3) |
| Completeness (%) | 99.6 (99.6) |
| $I / \sigma(I)$ | 6.9 (2.0) |
| $R_{\text{meas}}$ | 0.228 (1.246) |
| $CC_{1/2}$ | 0.982 (0.646) |
| *Refinement* |  |
| Resolution ($\text{\AA}$) | 37.12–2.30 |
| No. of reflections | 8367 |
| No. of protein atoms | 1566 |
| No. of water molecules | 29 |
| $R_{\text{work}}$ | 0.274 |
| $R_{\text{free}}^1$ | 0.326 |
| RMSD bond length ($\text{\AA}$) | 0.012 |
| RMSD bond angles ($^\circ$) | 1.499 |
| Average B-factor ($\text{\AA}^2$) | 42.76 |
| Ramachandran plots |  |
| in favored region (%) | 98.3 |
| in allowed region (%) | 1.7 |
| in outlier region (%) | 0.0 |
| PDB accession code | 9UGR |

All numbers in parentheses represent the last outer shell statistics.
<sup>1</sup> $R_{\text{free}}$ is calculated for 5% of randomly selected reflections excluded from refinement.

**SI Table 2.** Summary of small-angle X-ray scattering (SAXS) results.

| Sample | $I(q \rightarrow 0)/c$<br>[a.u.] | $D_{\max}$<br>[nm] | $R_g$<br>[nm] | $M_w$<br>[kDa] | Calc. $M$ [kDa]<br>(Monomer) | Oligomeric<br>State [mer] |
| --- | --- | --- | --- | --- | --- | --- |
| Syn-F4_Link | 0.01350 | 10.5 | 2.50 | 23.8 | 25.5 | 1 |
| Syn-F4(K4T) | 0.01314 | 10.6 | 2.58 | 23.2 | 12.5 | 2 |
| Lysozyme | 0.00812 | 6.0 | 1.51 | 14.3 | 14.3 | 1 |

